# A unique HMG-box domain of mouse Maelstrom binds structured RNA but not double stranded DNA

**DOI:** 10.1101/014522

**Authors:** Pavol Genzor, Alex Bortvin

**Affiliations:** Department of Embryology, Carnegie Institution for Science, Baltimore, Maryland, United States of America; Department of Biology, Johns Hopkins University, Baltimore, Maryland, United States of America

## Abstract

Piwi-interacting piRNAs are a major and essential class of small RNAs in the animal germ cells with a prominent role in transposon control. Efficient piRNA biogenesis and function require a cohort of proteins conserved throughout the animal kingdom. Here we studied Maelstrom (MAEL), which is essential for piRNA biogenesis and germ cell differentiation in flies and mice. MAEL contains a high mobility group (HMG)-box domain and a Maelstrom-specific domain with a presumptive RNase H-fold. We employed a combination of sequence analyses, structural and biochemical approaches to evaluate and compare nucleic acid binding of mouse MAEL HMG-box to that of canonical HMG-box domain proteins (SRY and HMGB1a). MAEL HMG-box failed to bind double-stranded (ds)DNA but bound to structured RNA. We also identified important roles of a novel cluster of arginine residues in MAEL HMG-box in these interactions. Cumulatively, our results suggest that the MAEL HMG-box domain may contribute to MAEL function in selective processing of retrotransposon RNA into piRNAs. In this regard, a cellular role of MAEL HMG-box domain is reminiscent of that of HMGB1 as a sentinel of immunogenic nucleic acids in the innate immune response.

## Introduction

Integrity of the germ cell genome is central to sexual reproduction. Gamete development presents an ideal environment for the selfish propagation of transposable elements (TEs) such as LINE-1 (L1) [1-5]. In mammals, retrotransposon expression peaks during a period of genome-wide reprogramming of the embryonic germline but is subsequently extinguished in a sex-specific manner [6-11]. Retrotransposon dysregulation is associated with an accumulation of DNA damage, meiotic abnormalities, chromosome segregation errors and embryo lethality [12-16]. A prominent role in transposon control belongs to the piRNA pathway that operates through a conserved group of primarily germline-restricted factors, including Piwi proteins, Tudor domain containing proteins, and a large set of accessory proteins required for various aspects of piRNA biogenesis and function [17-22].

We focus on mouse Maelstrom (MAEL), arguably one of the most enigmatic proteins of the piRNA pathway because of diversity of biological roles that have been attributed to it [23-29]. Most consistently, however, MAEL was shown to be important for retrotransposon regulation in *Drosophila* and mice [14, 24, 26, 28, 30, 31]. The mouse MAEL is dynamically localized throughout the developing germ cell, most prominently seen in cytoplasmic piP-bodies of fetal germ cells and chromatoid bodies of round spermatids, both of which are thought to be sites of transposon RNA processing [24, 30]. In the absence of MAEL, piRNA biogenesis is significantly perturbed causing infertility in male and female mice due to meiotic defects, an observation concordant with an elevated retrotransposon expression [14, 24, 28, 30]. MAEL-containing complexes from adult mouse testes are highly enriched for MIWI (one of three Piwi proteins of mice), Tudor domain-containing TDRD6, and processing intermediates of transposon and piRNA precursor RNAs [30].

MAEL protein possesses two domains - an amino-terminal high mobility group (HMG)-box domain and a MAEL-specific domain (MSD) with a presumptive RNAse-H-like fold [32, 33]. We examine MAEL HMG-box domain that has been shown to be important for function of Mael in *Drosophila* [26]. HMG-box domains are well known for interacting with DNA in either sequence-specific (e.g. transcription factors, such as SRY and SOX) or non-sequence-specific (e.g. structural chromatin proteins such as HMGBs) manners [35-41]. Their early identification led to extensive characterization of their peptide sequences, tertiary folds, nucleic acid binding capabilities, and other biophysical parameters [37, 42]. In addition to being appreciated for their DNA binding abilities, members of the HMG-box domain superfamily, such as HMGB1, can also bind RNA, and appear to play an important role in the innate immune response to immunogenic foreign nucleic acids [43-45].

Here we take advantage of the available H-NMR structure of human MAEL HMG-box domain (PDBID: 2cto) [34] to describe *in vitro* biochemical functions of its mouse counterpart. Our results indicate that the MAEL HMG-box domain shares sequence and structure characteristics as well as phylogenetic proximity to domain A of HMGB proteins. Nevertheless, we identify additional features of the MAEL HMG-box within each category that set it apart from non-sequence-specific (NSS) HMGBs as well as sequence-specific (SS) HMG-box proteins Specifically, we show that the MAEL HMG-box domain is not able to bind dsDNA and exhibits preference for structured RNA substrates *in vitro*.

## Material and Methods

### Cloning and Mutagenesis

Mouse Maelstrom cDNA was previously generated in the lab [28]. HMGB1a cDNA was obtained from Open Biosystems (clone 30849071). SRY HMG was PCR amplified from mouse testis 129S4 cDNA. *Drosophila melanogaster* maelstrom was amplified from *Drosophila* testis cDNA (gift of the X. Chen lab, Johns Hopkins University). All HMG domains (nucleotides 2-258 coding for first 85 residues) were amplified with Phusion polymerase (NEB) using primers listed in Table S1 in Tables. PCR products were sub-cloned into pGex6P2 expression vector (GE) between BamH1 and Not1 restriction sites. Selected residues were mutated using round-the-horn site-directed mutagenesis [46] and confirmed by sequencing.

### Protein expression and purification

Domains sub-cloned into pGex6P2 vector were transfected into BL21-DE3* cells (Life Technologies). Single clones were expanded overnight and inoculated into a large volume of terrific broth (TB) media supplemented with appropriate antibiotics. When the OD_600_ reached 0.6-0.8, the culture was moved to 18°C incubator and protein expression was induced with 250 mM IPTG for 12-16 hours. The cells were collected following day by centrifugation, washed once with 1xPBS and resuspended in lysis buffer (1x PBS, 5% glycerol, 1mM PMSF, 1mM TCEP, 1mM MgCl_2_, protease inhibitors (Pierce)). The cell suspension was supplemented with Lysozyme (Sigma) to the final concentration of 1-2 μg/ml and incubated on ice for 30 minutes with occasional mixing. Lysed material was then sonicated (4 repeats, 20 second sonication, 50% duty, Misonix 3000) with one-minute incubation on ice between repeats. The sonicated mixture was spun (4°C, SS-34 rotor, 30 minutes, 18000 rpm) and the supernatant carefully moved to syringe and filtered with Millex HV filters (Millipore) to remove contaminants. The GST-fusion protein was purified by gravity on the glutathione agarose resin (Sigma) at 4°C unless otherwise noted. The filtered lysate was bound to the glutathione resin, washed with 5 column volumes of low salt buffer (LSB: 1xPBS, 5% glycerol), 5 column volumes of high salt buffer (HSB: 1xPBS, 5% glycerol, 1M NaCl) and again with 2 column volumes of LSB. The protein was eluted with 10 mM reduced glutathione in LSB (pH ∼8.5). To remove the GST tag, the eluate was supplemented with 1mM EDTA, 1mM TCEP, PreScission protease (GE) and incubated at 4°C for 12-16 hours. Phospho-cellulose (PC) columns (2.5 ml) were prepared from dry PC resin (Whatman P-11) following procedures provided by Lorsch Lab (NIGMS). Briefly, 0.8g of the resin was stirred into 125ml of 0.5N NaOH for 5 minutes. After that the resin was washed with water until pH < 11 at which point 125 ml of 0.5N HCl was added and the solution was stirred for 5 minutes again. The mixture was then washed with water until pH > 4, at which point the resin was poured into disposable columns and equilibrated with desired buffer until pH_IN_ = pH_OUT_. The PreScission digested glutathione column eluate was diluted with B0 buffer (B0: 20mM Hepes pH 7.4, 10% glycerol, 2 mM DTT, 0.1 mM EDTA) to lower the salt below 75 mM. The diluted digest was then loaded onto 2.5 ml PC columns and allowed to bind by gravity. The column was washed with 5 column volumes of B100 buffer (B0 + 100mM NaCl). The protein was eluted with B500 buffer (B0 + 0.5M NaCl). The fractions with A_280_ > 0.05 were pooled and their buffer exchanged (PD-10 desalting columns) to storage buffer (SB: 1x PBS, 5% glycerol, 1 mM TCEP). To remove residual PreScission protease and un-cleaved protein, the eluate was passed over 0.5 ml glutathione/0.5 ml Ni-NTA column. The final eluate was concentrated using Vivaspin 6 centrifugal concentrators (Satorius) with 3 MWCO to desired protein concentration (>1 mg/ml). The concentration was measured at A_280_ in 6M Gnd-HCl, 20 mM Sodium Phosphate pH7.5 using calculated extinction coefficient and molecular weight (http://www.expasy.org) on NanoDrop 2000c. The final protein was aliquoted, flash frozen and stored at -80°C until use.

### Circular dichroism (CD) spectroscopy

CD measurements were collected on Aviv 420 instrument (Aviv Biomedical). Far-UV spectra were collected in 0.1 cm cuvette at 25°C. All proteins were 94-residues long at 0.1mg/ml concentration. The Samples were in 1x PBS, 5% glycerol at room temperature. The data were processed in Numbers (iWork, Apple Inc.) as previously described [47].

### Simple substrate preparation

The DNA oligonucleotides for each substrate were purchased desalted without purification (Operon) (Table S2 in Tables). The RNA oligonucleotides were ordered desalted with HPLC purification (Sigma Proligo) or prepared by *in vitro* transcription of PCR products with T7 promoter using HiScribe T7 high yield RNA synthesis kit (NEB) following manufacturers protocol (Tables S3, S4, S5 in Tables). Briefly, the synthesized RNA was purified with acid phenol:chloroform and precipitated with isopropanol. The precipitate was diluted in 1x CutSmart™ buffer (NEB), supplemented with RNAseIN inhibitors (Ambion) and de-phosphorylated with alkaline phosphatase (NEB). The precipitation was repeated. The precipitate was then purified on TBE-UREA polyacrylamide gels (Live Technologies) and the RNA was purified following crush-and-soak method [48]. Briefly, the appropriate size band was excised from the gel, crushed in the presence of PAGE elution buffer (0.3M NaOAc, 10 mM Tris 8.0, 1 mM EDTA pH 8.0) and frozen at -80°C for 30 minutes. The RNA was eluted by shaking the mixture overnight at 37°C and precipitated with isopropanol. The molar concentration was calculated based on the A_280_ readings in 10 mM Tris pH 8.0. The RNA was stored at -80°C until use.

### Complex substrate preparation

The oligonucleotides were designed with sufficient overlap and homology to specifically anneal. To create a four-way junction, 10 μl (100 μM) of each oligonucleotide was mixed in the annealing buffer (1x: 70 mM Tris pH 7.5, 10 mM MgCl_2_, 100 mM NaCl) to a final volume of 200 μl. The mixture was incubated in 95°C water bath for 5 minutes and allowed to slowly cool to room temperature. The annealed substrate was precipitated with EtOH (DNA) or isopropanol (RNA). Approximately 10 μg of annealed substrate was diluted in the binding reaction without protein (1x: 10 mM Potassium Phosphate pH 7.5, 50 mM KCl, 5% glycerol, 1 mM TCEP, 2.5 mM MgCl_2_), loaded into single lane of 12% Native page gel and ran at 105V for 1-2 hours. Lower concentration of the acrylamide was used for the RNA substrates > 75 bases. The bands were visualized using short wavelength UV shadowing and appropriate bands were excised and purified following crush-and-soak method described earlier. The molar concentration of each substrate was calculated using molecular weight and A_280_ readings. The oligonucleotides were aliquoted at desired concentration and stored at - 80°C until used. All double stranded (RNA, DNA) substrates were annealed and purified in the same fashion. The RNA oligonucleotides for hairpin substrates were ordered HPLC-purified (Sigma Proligo).

### RNA substrate structural considerations

To simplify interpretations, all the substrates made from ssRNA were designed with potential secondary structural characteristics in mind. The sequences were submitted to the Mfold server [49], using standard settings to identify thermodynamically favorable confirmations. All structures with negative free energy (ΔG) were considered as likely within the ensemble of tested RNA. The structures with +ΔG were considered as unlikely. This is based on the fact that base pairing provides -ΔG to RNA molecule allowing for spontaneous folding and secondary structure formation [50]. Therefore, in Mfold analysis, sequences that produce structures with only +ΔG are considered single-stranded, whereas an ideal hairpin sequence would produce only single structure with large -ΔG. The Table S3 in Tables contains the free energies and structures of the tested substrates identified by Mfold.

### Gel shift assays

The substrates were diluted to desired concentrations and end-labeled with γ-P^32^ using PNK (NEB). To account for number of ends, 5 μM of the hot ATP were used per 1 μM of DNA four-way junction. Unincorporated label was removed on P30 columns (Bio-Rad). To control for the loss of the shorter substrates on the P30 column, multiple substrates were labeled at the same time and their concentrations were normalized to the DNA 4WJ (largest substrate) using relative incorporated scintillation counts. Such prepared substrates were stored at 4°C until use, unless folding was required. To fold, the RNA substrates were supplemented with salts (50 mM NaCl, 2.5 mM MgCl_2_) and heated to either 55°C (< 50 bases) or 95°C (>50 bases) for 3 minutes and allowed to slowly cool to RT. The folded RNA was stored at 4°C until use. The protein was thawed on ice and then serially diluted to desired concentrations in water. The binding reaction was assembled by mixing the protein in binding reaction consisting of (1x) 10 mM Potassium Phosphate 7.5, 50 mM KCl, 5% glycerol, 1 mM TCEP, 0.1 mg/ml BSA, 2.5 mM MgCl_2_. The labeled substrate was added last to ∼1nM concentration in 10 μl final volume. The reaction was then incubated at room temperature for 30-60 minutes to equilibrate. The 12% native polyacrylamide (29:1), 1 mm thick TBE (1x) minigels gels were pre-run wit 0.5x TBE running buffer for 30 minutes at 105V in ice water-bath.

The wells were briefly rinsed, and 5 μl of the binding reaction was then carefully loaded onto running gels. The gels were run at constant 105V for long enough (1-4 hours) to achieve sufficient complex separation. At the end of the run, gels were extracted, rinsed, dried onto 3 mm Whatman paper at 80°C for 90 minutes, and exposed to storage phosphor screen for 12-24 hours. The image was acquired using Storm 860 molecular imager (Molecular Dynamics) with 100-micron resolution. The large RNA substrates were treated the same way but the complexes were resolved on large 6% native gels. In the competition experiments, the binding reactions were setup in the same manner as above but with protein concentration held constant and sufficient to achieve between 60-90% binding. Serially diluted unlabeled (cold) substrate was added up to 1 uM final concentration prior to addition of the radioactively labeled (hot) substrate.

### Data analysis

The images obtained from the Storm 860 were analysed in FIJI (GPL). The region of the gel was extracted, the pixels inverted onto black background, and background subtracted uniformly amongst all images. For each lane the region free and the region bound were selected using gel analysis feature and the area under the curve quantified using wand tool. Multiple complexes were all included in the region bound. The fraction bound was calculated using equation (1) and data plotted as the fraction bound versus protein concentration using Prism6 software (GraphPad). To calculate dissociation constant (K_D_), data was fit to modified Hill equation (2). The cold competition data was plotted as the fraction of bound hot substrate versus the concentration of the cold substrate using equation (3) and the dissociation constant of competitor (K_C_) was calculated with equation (4). All parameters in equations (2,3,4) were described previously [51].

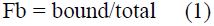

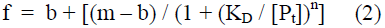

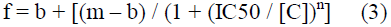

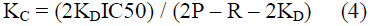

## HMG alignment and modeling

The full-length mouse Maelstrom sequence (434 residues) was submitted for tertiary structure prediction to the Robetta online server [52]. The .pdb files were retrieved and analyzed in PyMOL. The same process was followed for the *Drosophila melanogaster* Maelstrom HMG-box domain (residues 1-86). The .pdb files examined are provided in supplement.

Nucleotide sequences of the candidate sequence-specific (SRY, SOX) and nonsequence-specific (HMGB, Dsp1) HMG domains were obtained from NCBI and 86 residue region encompassing HMG box was selected for the alignment. The sequence id indicates protein name + species + start residue + number of consecutive residues extracted. The accession numbers used were: Maelstroms –[*Mus musculus* (Mm) Maelstrom - NM_175296.4, *Drosophila melanogaster* (Dm) Maelstrom - NM_079493.4, *Homo sapiens* (Hs) Maelstrom - DQ076156.2]; sequence-specific HMG proteins [Hs SRY - X53772.1, Mm SRY - NM_011564.1, Mm Sox2 - NM_011443.3, Mm Sox6 - U32614.1, Mm Sox10 - AF047043.1, Mm Sox17 - NM_011441.4]; non-sequence-specific HMG proteins [Dm Dsp1 - U13881.1, Mm HMGB1 - NM_010439.3, Mm HMGB2 - NM_008252.3, Mm HMGB3 - NM_008253.3]. Codon alignment was performed using the ClustalW algorithm built-in MEGA6 package, without changing pre-set parameters. The aligned nucleotides were translated to protein using standard genetic code and the alignment of protein repeated using built in ClustalW algorithm. No changes to codon alignment occurred. This alignment was manually refined using experimentally determined structural information to account for the secondary characteristics such as helices and loops. Following PDB structures were used: SRY – 1j46 [37], HMGB1a -1ckt [42], MAEL HMG – 2cto [34]. The final alignment was exported and the residues colored according to Taylor color scheme to reflect biochemical characteristics of various residues [53]. The final alignment along with annotation of the secondary structural elements is shown in Figure S1A. The alignment was then used to generate maximum likelihood tree using MEGA6 [54, 55] built-in algorithms with the following settings: 1000 Bootstrap replicates, Jones-Taylor-Thornton model of amino acid substitutions, uniform site-rates, complete deletion of gaps and missing data, Subtree-Pruning-Regrafting – Extensive at level 5, very strong branch swap filter. The generated tree was visually adjusted in built-in tree editor and is presented in Figure 1C. The log likelihood of this tree is -1943.6 and each branch is annotated with the bootstrap values representing the percentage of trees where the associated sequence clustered together. The tree branch scale represents number of substitution per site based on the considered 73 completely conserved positions amongst 17 compared sequences.

**Figure 1.**
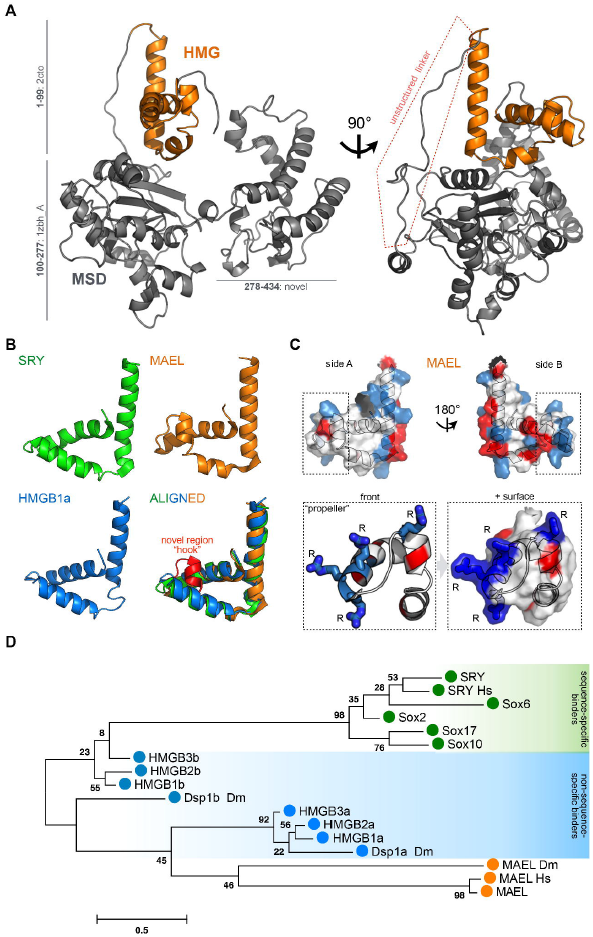
MAEL HMG-box domain structural and sequence considerations. (A) Tertiary structure prediction of mouse MAEL protein. MAEL HMG-box domain (orange) is positioned on top of the Maelstrom Specific Domain (MSD, grey) linked by unstructured linker (highlighted in red). The model was generated with Robetta software by either *de novo* modeling or with established structures as templates. Residue ranges and parent PDB ID numbers are shown. (B) Comparison of MAEL HMG-box and canonical HMG-box domains. Determined structures of candidate HMG-box proteins (SRY: 1j46 - sequence specific binding; HMGB1a: lckt - structure specific binding; MAEL: 2cto - unknown binding) were visualized and structures aligned in PyMOL. MAEL HMG-box domain has a conserved canonical L-shape fold but with a bend in helix-2, creating a novel region termed “hook” (red). (C) Distribution of charged residues of mouse MAEL HMG-box domain. Positive residues - Arg, Lys, His are blue; negative residues - Asp, Glu are red. Charged residues are concentrated on side B. Unlike other HMG-box domains, in MAEL three arginine (R) residues are concentrated at the end of the helix-1, and protrude outwards, forming a “propeller”-like shape. (D) Phylogenic comparison of MAEL HMG-box domain with well-studied candidate HMG-box domains from sequence-specific (single HMG-box, SRY, Sox) and non-sequence-specific (two HMG-boxes, HMGB’s, Dspl) groups. Mouse sequences were used unless otherwise noted (Dm: *Drosophila melanogaster*, Hs: *Homo sapiens)*. The phylogenetic tree was generated using maximum likelihood method in MEGA6 software. Values next to the branches describe percentage of trees where associated sequences group together (n=lGGG). The branch length scale is in substitutions per site. MAEL HMG-box domain forms a new branch most closely related to the domain A of nonsequence specific HMG-box proteins.

### Large RNA structure determination

Previously described MAEL RIP-Seq data sets mapped to mm9 assembly of the mouse genome, shown to be enriched in transposon RNA, were used for the identification of overrepresented regions [30]. The sets corresponding to control Igg, MAEL_A RIP and MAEL_B were analyzed with macs software (version 1.4.2) [56] with the standard settings to identify regions enriched in replicates A and B over Igg. The identified regions between the two replicates were pooled and intersected using bedtools (v2.20.1) [57] to identify only the common regions. All intervals were then annotated using annotatePeaks program from HOMER suite [58]. The regions annotated as LINE1 elements were extracted and their coordinates examined in IGV (Broad Institute), considering only the regions within annotated LINE1 elements. Multiple coordinates corresponding to regions with a peak appearance at least 250 nucleotides-wide were selected, and their nucleotide sequences extracted from the UCSC genome browser. These were then aligned using ClustalW (EMBL-EBI) and the alignment manually curated until the region of high sequence conservation was identified. The final alignment had 5 regions corresponding to LINE1 elements of Md_F2 family that were located on different chromosomes (Figure S5A). Coverage across each identified region was calculated using its coordinates and the bedtools multicov program [57]. The results were plotted in Numbers (iWork, Apple Inc.) (Figure S5B). This alignment was used for determination of the secondary structure according to previously described methodology [59]. The covarying nucleotides used to constrain Mfold [49] are provided in Table S6 in Tables. The region with lowest dG (chr10) was tested in gel shift assays.

## Results

### Structural overview of Maelstrom

To gain insights into the function of mouse Maelstrom (MAEL) protein, we first used Robetta protein structure prediction server [52] to predict its tertiary structure. The server utilizes sequence homology with previously determined structures as parents for the structure prediction. *De novo* methods are used if these are not available. In accordance with Maelstrom gene annotation (UniProt) and previous analysis [32, 33], we have annotated the resulting structure with two domains: an N-terminal HMG-box and a MAEL-specific domain (MSD) (Figure 1A). The predicted structure of mouse MAEL HMG-box domain is based on a previously obtained H-NMR structure of HMG-box domain of human MAEL protein (PDBID: 2cto) [34]. The MSD domain has been previously computationally predicted to assume an RNase H-like fold [32, 33]. In agreement with these studies, Robetta utilized an exonuclease structure (PDBID: 1zbh – chain A) as a parent molecule for this domain. The C-terminal sequence of MAEL protein was modeled *de novo* as it appears unique. The predicted structure shows the HMG-box domain on the surface and not encapsulated by the rest of the molecule. Instead, it is connected with the MSD domain by an approximately 30-residue linker region that appears devoid of any secondary structural elements (Figure 1A). Based on the sequence composition, this linker region is predicted to have high propensity for intrinsic disorder, which could account for insolubility that we have encountered while attempting to purify recombinant full-length or truncated Maelstrom proteins. The fact that the MAEL HMG-box domain is not buried, but instead connected to rest of the protein with an unstructured linker reaffirmed our interest in understanding its function.

### The HMG-box domain of Maelstrom

MAEL is the only known HMG-box domain-containing protein in the piRNA pathway. Structurally, all HMG-box domains have a characteristic L-shape fold of three helices (Figure 1B) [40]. Like the HMG-box domains of SRY and HMGB1 proteins, mouse MAEL HMG-box domain also possesses this basic fold. However, it has acquired novel features, most prominently a distinguishable bend in helix-2 apparent from a simple structural alignment (Figure 1B). This change of geometry gives helix-2 the appearance of a "hook". Because the equivalent region is known to be important for binding of canonical HMG-box domains [40], we predict this "hook" region to have functional consequences for the mouse MAEL HMG-box domain. Surface rendering of this domain shows that the region is bulky, containing three consecutive arginines at the C-terminus of the helix-1 that form what appears to look like a “propeller” (Figure 1C). The consecutive positively charged residues are present in the canonical HMG-box domains (Figure S1). However, in sequence-specific (SS) binders these residues are located internally in helix-1, while in non-sequence-specific (NSS) binders these are not arginines (Figure S1). The presence of arginines is significant due to their ability to form multiple H-bonds with nucleic acid bases or the backbone [60, 61]. In addition, arginine residues may also be involved in recognition of specific motifs within RNA [62]. Importantly, the “hook” and the “propeller” are specific to mouse MAEL HMG-box, and could be of functional significance.

To infer the domain relationships, we performed multiple sequence alignment of candidate HMG-box domains from SS and NSS groups with the mouse, human and *Drosophila* Mael HMG-box domains (Figure S1). This analysis showed that the MAEL HMG-box domains form a separate branch on the phylogenic tree (Figure 1D). In addition to the described structural differences specific to the mammalian MAEL HMG-box domain, this implies that there are other features common to the MAEL HMG-box domain homologues that may be important. The MAEL HMG-box domains are most closely related to domain A of non-sequence-specific binders, but differ from these in their distribution of charged residues (Figure S1). In the mammalian MAEL HMG-box domains, the loop connecting helices-1 and 2 does not contain charged residues, and helix-2 is devoid of the positively charged residues that are present in all other groups (Figure S1). While charged residues in other domains seem to be alternating from helix-1 to helix-2, the mammalian MAEL HMG-box domain has concentrated positive residues, which form a novel region. The distribution of charged residues is indicative of an H-bond potential that, together with non-polar regions, can provide the biochemical basis for strong interactions with appropriate substrates. These features vary in *Drosophila* Mael HMG-box domain that is still distinct from canonical HMG-boxes (Figure 1C, Figure S1A). The HMG-box domain of the *Drosophila* Maelstrom protein has evolved distinct features from its mammalian homologues, which perhaps reflect specie-specific specialization required for its functions. Nevertheless, all analyzed MAEL HMG-box domains, while related to the canonical HMG-box domains, have evolved characteristics that set them apart, and these are likely to influence their function.

### Binding of MAEL HMG-box domain to DNA

To evaluate the biochemical activity and validate our previous analysis of MAEL HMG-box domain, we expressed HMG-box domains of SRY (SS), HMGB1a (NSS) and murine, and *Drosophila* Mael HMG-box domains in bacteria (Figure S2A). The recombinant proteins were purified to homogeneity and then CD analysis was used to confirm their tertiary structure (Figure S2B-D). To evaluate the ability of HMG-box domains to bind nucleic acids, we used gel shift assays where we titrated increasing amounts of protein of interest to known concentration of labeled (hot) substrate and determined the dissociation constant (K_D_). When appropriate, we utilized competition assays, where the protein concentration was kept constant (60-90% total binding) along with the concentration of the hot substrate, and instead the unlabeled (cold) substrate was titrated in. This allowed us to estimate the competitor dissociation constant (K_C_) that is directly related to the dissociation constant obtained from the binding assay. Considered together these two constants provide an estimate of the observed binding kinetics obtained by gel shifts.

We first evaluated DNA binding of SRY HMG-box and HMGB1a recombinant proteins (Figure 2A). Previous studies have determined that the SRY HMG-box domain binds dsDNA in a sequence-specific manner with consensus sequence AACAAN [35]. The SRY HMG-box domain recognizes the sequence through a number of minor groove interactions, intercalates a residue at the beginning of its helix-1 between the bases, bending the helical backbone and allowing for accommodation of rest of the helix in the minor groove [37]. Due to its specific residue composition, the SRY HMG-box domain is able to bend the DNA. Consistently, recombinant SRY HMG-box domain bound its consensus sequence strongly (average K_D_ = ∼12 nM) and specifically forming a single complex (Figure 2A, Figure S3A). In contrast, HMGB1a is known to bind pre-bent but not unperturbed dsDNA as it lacks the residues required for DNA bending [39, 40, 42]. This is in accordance with our observation that HMGB1a does not bind same dsDNA (Figure 2B).

**Figure 2.**
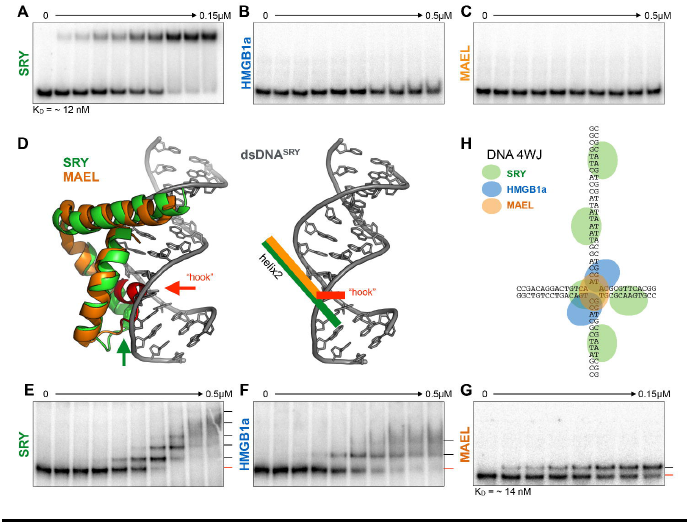
MAEL HMG-box domain binding to DNA. (A) Recombinant SRY HMG-box domain strongly binds to dsDNA with its consensus sequence (dsDNA^SRY^). (B) Recombinant HMGB1a protein does not bind to the same substrate because it is unable to bend linear dsDNA. (C) Similar to (B) MAEL HMG-box does not bind to dsDNA. (D) Representation of co-crystal structure of SRY HMG-box (green) with dsDNA^SRY^ describing their fit. Aligned with SRY HMG-box is the MAEL HMG-box domain (orange) whose “hook” region protrudes deeply into dsDNA (1j46, 2cto). The geometry of the helix-2 of both molecules in relation to dsDNA^SRY^ is highlighted on the right. Even though dsDNA^SRY^ is pre-bent, it cannot accommodate the “hook” region of MAEL HMG-box domain in the same fashion. (E) SRY HMG-box domain binds to DNA 4WJ forming five complexes (red lines - free substrate; black line - bound substrate). (F) HMGB1a binds to DNA 4WJ forming two complexes. (G) The MAEL HMG-box domain forms only single complex with DNA 4WJ, which is different from the SRY and HMGB1a domains. (H) Proposed mode of HMG-box domain binding to DNA 4WJ. SRY HMG-box domain recognizes a distorted 4WJ center as well any sequences that approximate its consensus-binding site, while HMGB1a protein binds primarily to irregular center. Just as HMGB1a, MAEL HMG-boxes does not bind dsDNA suggesting that it also binds to the center of the junction. Because only single complex forms binding likely happens in different fashion.

The MAEL HMG-box domains do not bind to single stranded (ss) (Figure S3B - mouse), dsDNA (Figure 2C - mouse, Figure S3C - *Drosophila)*, or dsDNA methylated at CpGs (Figure S3D - mouse). We have tested the mouse and *Drosophila* MAEL HMG-boxes with multiple dsDNA substrates containing canonical HMG-box motifs and non-canonical sequences, however we failed to detect complex formation under our conditions (Table S2 in Tables). Likely the reason for lack of mouse MAEL HMG-box binding is the "hook" region that prevents accommodation of the protein helices in the dsDNA groves even when the dsDNA is modified or pre-bent (Figure 2D). Even though sequence and predicted structure of the *Drosophila* Mael HMG-box do not show a homologous “hook” region, it is still unable to bind dsDNA (Figure S1, Figure S3D). The presence of two arginines in the *Drosophila* domain suggests that an analogous feature may also be present (Figure S1). We have not tested all possible sequences for binding, however additional sequence permutations would only produce dsDNA with the B-type helix. Therefore, it is highly probable that the structural characteristics of the MAEL HMG-box domain will prevent any significant interactions in a sequence-specific manner.

A common observed characteristic of the HMG-box domains is the ability to bind to DNA four-way junctions (4WJ) in their open conformation [41, 63, 64]. These junctions are comprised of four double stranded arms with a central junction where the strands sharply turn and the helical grooves widen, essentially providing a pre-bent and open site for binding. To determine whether mouse MAEL HMG-box domain has retained this characteristic, we have compared its binding with that of SRY HMG-box and HMGB1a, both of which bound DNA 4WJ, to readily form multiple complexes (Figure 2E-F). SRY formed five complexes with DNA 4WJ while HMGB1a formed two. However, MAEL HMG-box domain was able to form only a single complex even at high protein concentrations (Figure 2G, Figure S3D). Binding of SRY and other HMG-box domains to DNA 4WJ, as well as dynamics and structure of DNA 4WJ are well described [41, 63-78]. In accordance with these previous observations, the five SRY-DNA 4WJ complexes are likely the products of binding of SRY HMG-box domains to the 4WJ open center and to the AT-rich sites in double-stranded arms that approximate SRY recognition sequence (Figure 2H). The two HMGB1a complexes likely represent two protein domains symmetrically accommodated at the irregular center of the junction. Only a pre-bent center can be bound due to the lack of intercalating residues required for bending of and consecutive binding to unperturbed dsDNA by HMGB1a [42, 68, 79]. Like HMGB1a, the MAEL HMG-box domain does not bind to dsDNA, but unlike HMGB1a, only a single complex is formed with the DNA 4WJ (Figure 2H). A possible explanation for this is that the “hook” and “propeller” regions are accommodated at the open center of the junction, however, their bulkiness prevents accommodation of the second protein.

Taken together, the above experiments show that MAEL HMG-box domain does not bind to dsDNA, however it is able to form a single strong and specific complex with DNA 4WJ (K_D_ = ∼l4 nM, Figure S3E). This mode of binding is different from that of tested canonical HMG-box domains (Figure 2E-F). Considering the unique structural characteristics of MAEL HMG-box domain, we propose the “hook” and “propeller” regions may play an important role in what appears to be a not sequence but a structure-specific mode of binding.

### Binding of MAEL HMG-box domain to RNA

The MAEL is an important member of the piRNA pathway and specifically immunoprecipitates with piRNA precursor transcripts and transposon mRNAs [30]. Furthermore, the presence of RNAseH-like domain suggests that MAEL may operate in the RNA context [32, 33]. Considered together with herein observed binding to structured DNA, we wanted to probe MAEL HMG-box binding to RNA.

The cellular RNAs exist as single-stranded molecules that are capable of forming intricate secondary and tertiary structures [50]. Therefore, using Mfold, we have identified conformational ensembles of RNAs to be tested for MAEL HMG-box binding (see Methods, Table S3). While we did not observe any binding of the MAEL HMG-box domain to ssRNA (Figure 3A), we did detect a weak complex formation with dsRNA (Figure 3B). The MAEL HMG-box domain did not bind to small hairpin structures (Figure 3C-D), but formed weak complexes with larger RNA hairpins (Figure 3E-G). Of these, the strongest binding was observed with the RNA hairpin that carried the longest continuous dsRNA stem (9 base pairs) and hairpin loop (7 bases) (Figure 3F). Because only ∼40% of this substrate was bound at a relatively high protein concentration, we were not able to calculate binding parameters. Lastly, we tested the RNA counterpart of the DNA 4WJ junction used previously (Figure 2G, H) of the same nucleotide sequence but with RNA bases (Table S3). MAEL HMG-box domain bound well to this substrate with single complex forming at lowest tested concentrations and multiple complexes formed at highest concentrations (Figure 3H). Even though complete binding was not achieved, the binding strength was moderate (K_D_ = 0.638 μM, Figure S4A). The first complex formation resembled the interaction observed with DNA 4WJ, but never reached completion. Formation of the large complex was not observed previously with DNA 4WJ (Figure 2G). In order to get at the specificity of the binding, we attempted a competition assay with the cold substrate, but the results were rather puzzling. Instead of cold substrate titrating away the protein from hot substrate, all of the hot substrate shifted to large complex (Figure S4B).

**Figure 3.**
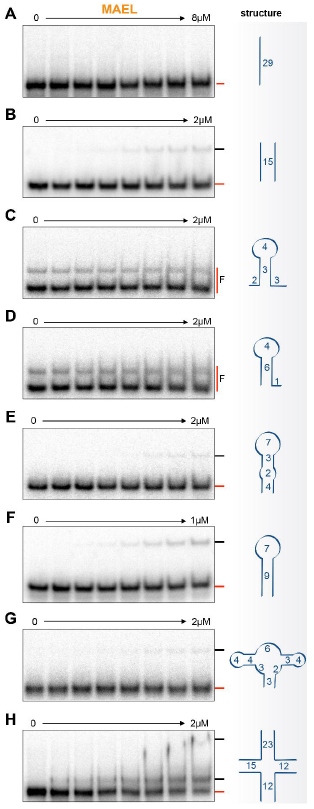
MAEL HMG-box domain binding to small RNAs. A) MAEL HMG-box domain does not bind to ssRNA. B) However, it forms a complex with double-stranded (ds) RNA of identical sequence as dsDNA^SRY^. C, D) MAEL HMG-box does not bind to small RNA hairpins. E) Only weak complex formation is observed with hairpin that has mismatches in the stem. F) But when the stem is perfectly base-paired, the binding is stronger and appears increased than that observed with dsRNA (B). G) Binding of MAEL HMG-box to substrate with multiple short hairpins is weak. H) However, binding to RNA 4WJ (sequence identical to DNA 4WJ) is strong and two complexes are formed. These observations indicate that MAEL HMG-box prefers to bind to substrates with continuous dsRNA helices longer than 6 base-pairs located near unstructured or perturbed RNA regions.

The above results suggest that MAEL HMG-box domain prefers RNA hairpins with completely base-paired stems with adjacent loops larger than 4 bases to ones with disrupted double-stranded regions and smaller hairpins. MAEL HMG-box domain interacts better with RNA 4WJ than with other RNA tested earlier. However, unlike with DNA 4WJ, it forms a large complex (Figure 3H). Furthermore, instead of being titrated away, this complex became predominant with additional RNA in the reaction (Figure S4B). A description of identical RNA 4WJ by others [80] suggests a possible explanation for the presence of these larger complexes. Unlike a DNA 4WJ that primarily exists in the open conformation [63], the RNA counterpart is more dynamic, undergoing multiple structural transitions [63, 80]. Therefore, its ensemble is largely composed of structures with dsRNA arms that are adjacent to each other either in parallel or antiparallel orientations (Figure S4C). It is thus possible that RNA helices in proximity to each other form a structurally unique region that accommodates multiple MAEL HMG-box domains at once. This mode of binding is supported by the arginine-rich sequence and the structure of MAEL HMG-box domain (Figure 1, S1). These positive residues in the "hook" and the "propeller" regions are distributed such that they span almost 270 degrees, providing sufficient rotational freedom for the rest of the domain to be accommodated in multiple ways. Arginine-rich peptides are enriched in other RNA-binding motifs and, have previously been implicated in facilitating complex protein-RNA interactions through “arginine-fork” phenomena [62, 81, 82]. Therefore, formation of large complexes with RNA 4WJ may be due to the interaction of arginine-rich regions of MAEL HMG-box domain with the closed portion of 4WJ ensemble. In the presence of additional RNA 4WJ in the reaction, the ensemble of the structures effectively changes to favor closed conformations (Figure S4C). HMGB1a binds RNA 4WJ similarly to its DNA counterpart, progressively forming larger complexes as more protein is bound (Figure S4D, Figure 2F). In comparison, MAEL HMG-box domain forms a single complex with DNA 4WJ. A similar complex is observed with RNA 4WJ, but an additional complex is observed without apparent intermediate states (Figure 2G, S3E, 3H, S4B). Additionally, the positive residues found within HMGB1a are all lysines, which are not capable of interactions equivalent to arginines despite their similar charge [62]. Taken together, the RNA binding data suggest that the MAEL HMG-box domain binds to RNA in a complex manner employing its arginine-rich “hook” and “propeller” regions to bind to structured RNA.

### MAEL HMG-box domain mutagenesis

In order to determine whether arginines in the “hook” and “propeller” regions of the MAEL HMG-box domain are important for binding, we have mutated the individual arginines to alanines. Additionally, we have mutated the glutamine (Q16) along the helix-1 and the arginine (R8) in N-terminus of the helix-1 to see whether polar residues within these regions are also important for binding (Figure 4A). Mutation Q16A in the middle of helix-1 had no effect on binding to the DNA 4WJ, however changed the complexes formed with RNA 4WJ (Figure 4B, B`). Differential binding to DNA versus RNA 4WJ could be due to differences in ensemble composition of the two junctions, which accommodate MAEL HMG-box domain in very different fashions despite identical sequences. The R8A mutation completely abolished binding to both DNA and RNA, indicating that this residue may be important (Figure 4C, C`). However, upon further inspection, the secondary structure of R8A was found affected (Figure S2B), therefore, the loss of binding might reflect changes in protein folding. In contrast, mutations R23A and R25A in the "propeller" region and R31A in the “hook” region had no effect on protein folding but completely abolished binding to DNA 4WJ (Figure 4D-F, Figure S2B). This result supports our previous hypothesis that arginines in this region are distributed such that they form multiple contacts with the perturbed region of this substrate. Binding of the same three mutants (R23A, R25A, R31A) to RNA 4WJ was significantly decreased (Figure 4D`-F`) compared to wild type (Figure 3H). Mutating "propeller" residues (R23A, R25A) allowed for the formation of a large complex, but mutation of the "hook" arginine (R31A) abolished it. Instead, a small amount of a new complex at an intermediate position was observed (Figure 4F`). These results also support our previous conclusions highlighting the importance of arginines in the "hook" and the "propeller" regions. It has been previously noted that even individual arginine residues are able to exert some degree of binding through “arginine-fork” phenomenon [62]. Therefore, considering the structural differences and the ensemble complexity of RNA 4WJ (versus DNA), the observed small amount of the complex in a single mutant binding assays may be a result of three arginines still being present and accommodated in one of many possible orientations.

**Figure 4.**
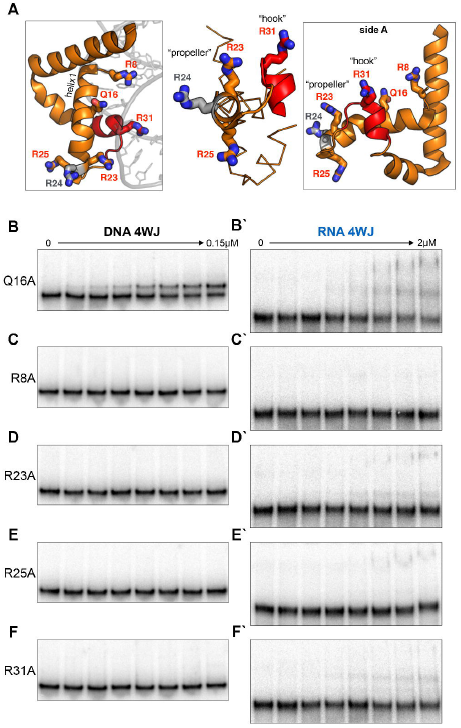
Site-specific mutagenesis of MAEL HMG-box domain. (A) Charged residues, capable of H-bond formation with long side chains, along helix-1 (region important for canonical HMG-box binding) and novel regions of MAEL HMG-box domain were mutated to alanine (residues in red). (B) Mutation of glutamine (Q16A) does not affect binding suggesting that this residue does not participate in complex formation with DNA 4WJ. (C-F) Mutation of individual arginines (R) results in the complete loss of binding to DNA 4WJ suggesting that they are essential for complex formation. The R8A mutant fold appears to be affected (Figure S2B), which might be responsible for reduced binding. Therefore, only arginines in the “hook” and “propeller” seem to be necessary for MAEL HMG-box domain binding to non-canonical DNA. (B’-F’) Interactions of mutants with RNA 4WJ are significantly affected. All mutants show greatly decreased binding to RNA 4WJ as well as variability in the type of complex formed when compared to wild-type protein (Figure 3H). Residual complex formation was still observed (except for R8A) likely due to a high number of configurations that RNA 4WJ can take to accommodate the MAEL HMG-box domain. Nevertheless, the arginines in the “hook” and “propeller” of MAEL HMG-box are important for binding to structured RNA.

The overall mutational analysis of the MAEL HMG-box domain has revealed that arginines in the "hook" and "propeller" regions are essential for binding, supporting our sequence and structure analyses and the interpretation of previous binding experiments. Taken together, our results point towards structured RNA as the preferred substrate for the MAEL HMG-box domain.

### MAEL HMG-box domain binding to large structured RNA

The apparent preference of MAEL HMG-box domain for structured RNA is in agreement with the results of our analysis of MAEL immunoprecipitates (IP). MAEL protein complexes immunoprecipitated from the adult mouse testis lysate are specifically enriched for the fragments of piRNA precursor RNAs and retrotransposon mRNAs [30]. Therefore, we explored the possibility that the MAEL HMG-box domain bound to endogenous long RNAs. We searched MAEL IP RNA-Seq data for enriched L1 sequences and identified a fragment of the L1_Md_F2 element (Figure S5A-B). Repeated and structured regions in mouse L1 elements have been previously described but no specific recognition signature has been defined [83, 84]. Under the assumption that retrotransposon L1 RNA can be under positive selective pressure to retain some structural features, we determined limited secondary structure of the identified regions of L1_Md_F2 using a combination of covariation and thermodynamic approaches. Such secondary structure would allow us to examine various features that may be bound by MAEL HMG-box domain. To do this, we followed methodology applied previously to determine the structure of yeast telomerase flexible scaffold [59]. Initial attempts to determine the structure of a 277-nt long piece of L1_Md_F2 with Mfold [49] produced multiple distinct structures making it impossible to identify their common structural features. After supplying the program with the identified covarying nucleotides, Mfold produced a single highly energetically stable structure for each region (Figure S5C). It was reassuring to see that the structures originating from sequences located on different chromosomes resembled each other in both sequence and structure, with only minor variations. For further analysis, we selected the most energetically stable structure corresponding to the sequence from chromosome 10 (Figure 5A). MAEL HMG-box domain bound this RNA substrate strongly, forming a single complex starting at 0.5 μM protein (Figure 5B). Interestingly, the mobility of the complex in the gel was progressively more retarded with increasing amount of protein in the reaction. The binding of MAEL HMG-box domain to this long RNAs does not occur with the same kinetics as with DNA 4WJ (Figure 2G), where a single domain binds to a single region in a non-cooperative manner. Instead, binding of long RNA appears to be highly cooperative, similar to the RNA 4WJ large complex (Figure 3H, Figure S4B), with consecutive molecules being bound after passing a certain concentration threshold of protein. In an attempt to identify the region that is recognized within this long RNA, we generated two shorter substrates, removing doublestranded segments of the stem, to create 200 and 149 nt long RNAs (Figure 5A). While the full-length fragment was completely bound, only fractions of the shorter substrates were shifted even at the highest concentrations of protein (Figure 5B).

**Figure 5.**
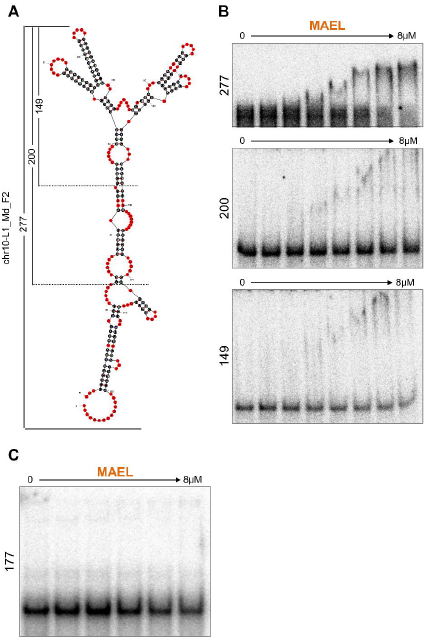
MAEL HMG-box domain binding to long RNAs. (A) The secondary structure of a conserved fragment of L1_Md_F2 element from chromosome 1O enriched in MAEL immunoprecipitates determined using combination of covariance and thermodynamic analyses. (B) MAEL HMG-box domain forms strong multiprotein complex with 277-nt fragment and binds weakly to shorter fragments (2OO-nt, 149-nt) of the same RNA. This suggests a requirement for complex tertiary structures in long RNAs for strong binding to the MAEL HMG-box domain. (C) MAEL HMG-box domain does not bind long RNAs originating from regions not enriched in MAEL immunoprecipitates.

These observations suggested that the removed stem somehow contributes to binding. Perhaps the double-stranded region constrains the ends of the RNA molecule allowing for unambiguous formation of the dual hairpin regions in the center. In this way, the ensemble of structures would be smaller with greater proportion of the preferred substrate. This would also explain the presence of the weak complexes and the lack of complete binding seen with shorter RNAs. Presence of additional RNA in reactions with RNA 4WJ have led to full formation of large complexes, most likely also affecting the ensemble of structures (Figure S4B). Importantly, testing the long RNA from a region adjacent to those recovered from MAEL immunoprecipitates failed to show any appreciable binding (Figure 5C). Our analysis of the single transposon RNA is limited to a single structured fragment and flanking region and therefore its implication should be considered with caution. Nevertheless, with the previous biochemical observations, it raises the possibility that the MAEL HMG-box domain contributes structure-specific RNA binding ability to the MAEL protein and in such way may aid in selection of MAEL-immunoprecipitated RNAs.

## Discussion

The aim of the study is to shed light onto the biochemical function of MAEL, a protein indispensable for the function of the piRNA pathway [24, 28, 30, 85]. We focused on the N-terminal HMG-box domain, which is important for MAEL biological function [26]. Its classification implies DNA binding ability, which is the case for many canonical HMG-box domains [35-37]. However, MAEL has been almost exclusively linked to the piRNA pathway where it is essential for piRNA biogenesis [24, 28, 30] and localizes to cytoplasmic piP-bodies and chromatoid bodies, likely involved in retrotransposon mRNA and piRNA precursor RNA processing [24, 30]. Additionally, retrotransposon RNAs are strongly enriched in MAEL immunoprecipitates [30]. Therefore, the evidence trail led us to hypothesize that the MAEL HMG-box domain is involved in RNA binding.

A plethora of non-sequence specific RNA binding domains have been described [86], but very few of them have the HMG-box domain [43, 44]. Our sequence and structure analysis of MAEL HMG-box domain indicates that it belongs to this exclusive group. The MAEL HMG-box domain does not bind single-stranded, double-stranded, or modified DNA molecules *in vitro* despite the presence of the consensus HMG-box binding sites in the tested substrates. Of the DNA substrates, MAEL HMG-box domain only binds DNA 4WJs, where it likely interacts with the structured center like many other HMG-box domains [63, 64]. On the other hand, it readily binds to RNA hairpins and forms multiple complexes with RNA 4WJ. As opposed to DNA, RNA junctions are far more prevalent in the cellular environment and commonly found in large molecules [87, 88]. Furthermore, we describe a case where, MAEL HMG-box domain is able to preferentially bind to a large structured RNA molecule originating from MAEL immunoprecipitates. Based on these observations, MAEL HMG-box domain could provide structure-specific RNA-binding capability to the full-length MAEL protein.

We have identified a region within MAEL HMG-box domain rich in arginine residues that is responsible for complex formation with the structured nucleic acid substrates.

The role of arginine-rich protein motifs in RNA binding have previously been demonstrated [82]. In MAEL HMG-box domain, the arginines residues form bulky "hook" and "propeller" regions providing a charged surface that, as our mutational studies showed, is required for strong and specific interactions with nucleic acids, likely through formation of arginine-forks [62]. The fact that even a single arginine mutation significantly affects the binding demonstrates that the composition and the architecture of these regions are important. Our work also suggests that MAEL HMG-box domain’s "hook" and "propeller" regions set it apart from known HMG-box domains, contributing to the formation of a phylogenetically distinct group of MAEL HMG-boxes. Given the exclusivity of MAEL HMG-box domain in the piRNA pathway, it is tempting to speculate that it has diverged and acquired the described features to accomplish a novel function perhaps specific to the piRNA pathway. Such function could involve discrimination of L1 and piRNA precursor RNAs from other transcripts. An *in vitro* preference of MAEL HMG-box domain for structured nucleic acids, including RNA hairpins, four way junctions, and large structured RNAs are all in agreement with this hypothesis. We believe that the combination of new *in vitro* (RNBS, [89]) with *in vivo* techniques (HITS-CLIP, [90]) in the future will reveal whether this hypothesis is correct.

Lastly, a biochemical activity of the MAEL HMG-box domain *in vitro* is reminiscent of that of HMGB1a in terms of structure-directed binding. Interestingly, in addition to their prominent structural role in the nucleus, HMGB proteins have been shown to function as sentinels of immunogenic nucleic acids in innate cellular response [44, 45]. A parallel presents itself where MAEL HMG-box may have diverged to aid in recognition of domesticated transposon RNAs. In this context, the piRNA pathway may be considered as an ancient arm of the innate immune response to protect genomes against retroviruses [91].

## Acknowledgements

We would like to thank Rejeanne Juste (Carnegie Institution of Washington) for performing cloning and mutagenesis; Johns Hopkins Center for Molecular Biophysics for help in acquisition of CD measurements; Xin Chen lab (Johns Hopkins University) for fly testis cDNA, Jon R. Lorsch (NIGMS) for help with PC column purification, Joseph-Kevin Igwe for help with HMGB1a purification; Safia Malki (Carnegie Institution of Washington), Valeryia Gaysinskaya (Carnegie Institution of Washington, Johns Hopkins University), Marla Tharp (Johns Hopkins University) for manuscript critique; Julio Castañeda (Carnegie Institution of Washington, Johns Hopkins University), David Zappulla (Johns Hopkins University), Alan Spradling (Carnegie Institution of Washington, Johns Hopkins University), Sarah Woodson (Johns Hopkins University), Chen-ming Fan and Christoph Lepper (Carnegie Institution of Washington) for critical discussion. Michelle Rozo (Carnegie Institution of Washington, Johns Hopkins University for help with manuscript revisions.

## Supporting Information Captions

**Figure S1. Sequence and secondary structural comparison of HMG-box domains.**

An amino acid multiple sequence alignment of candidate HMG-box domains. The candidates were selected based on their substrate specificity (sequence vs. non-sequence specific) with at least four candidates per group, and with preferences for well-described and mouse HMG-box domains. Aligned codons (ClustalW) were translated to protein sequences (MEGA6), and then the alignment was adjusted to account for the secondary structural characteristics found in solved structures within each group (SRY HMG-box -1j46, HMGB1a-1ckt, MAEL HMG-box -2cto). The residues were pseudo-colored according to Taylor color scheme (JalView) to provide contrast to groups of residues. Conserved secondary structure characteristics and residues are shown below the alignment.

**Figure S2. Purification of recombinant HMG-box domains.**

(A) Purification scheme employed for all recombinant HMG-box domain proteins. Shown are example gels demonstrating purity of acquired proteins. (B) Circular dichroism (CD) measurements of recombinant wild-type and point-mutant mouse MAEL HMG box domain proteins. All proteins are folded and helical. Only mutant R8A shows change in ellipticity (arrow) indicating that this mutation somehow affects folding of the protein. CD traces of C) *Drosophila* MAEL HMG-box, D) SRY HMG-box, and E) HMGB1a domains.

**Figure S3. HMG-box domain interactions with DNA substrates.**

(A) Binding of SRY HMG-box domain to dsDNA with its consensus recognition sequence (dsDNA^SRY^). Top gel shows the titration series for SRY HMG-box domain protein. The bottom gel shows cold competition with unlabeled dsDNA demonstrating strength and specificity of binding. The average K_D_ ∼12 nM is close to previously reported values [95]. (B-C) The mouse MAEL HMG-box does not bind to 25-nt ssDNA or dsDNA modified with symmetrical cytosine methylation. (D) Predicted (Robetta) tertiary structure of *Drosophila melanogaster* (Dm) Mael HMG-box domain. Based on the predicted structure, this domain closely resembles canonical HMG-box domains without any apparent novel regions. Despite the homology and contrary to previous observations dsDNA [27], we did not detect binding to dsDNA. (E) Binding of mouse MAEL HMG-box domain to DNA 4WJ. Top two gels show titration of protein with different concentration ranges. Even at high excess only single MAEL HMG-box domain is able to bind to DNA 4WJ. The bottom gel represents cold competition with unlabeled DNA 4WJ. The binding of the MAEL HMG-box domain is strong: average K_D_ ∼14 nM.

**Figure S4. MAEL HMG-box domain interactions with RNA substrates.**

(A) Strength of MAEL HMG-box domain interaction with RNA 4WJ is moderate (K_D_ = 0.638 μM). (B) Cold competition of MAEL HMG-box domain with RNA 4WJ leads to increased formation of the large complex instead of competition. Presence of more RNA likely influences structural ensemble, making the formation of large complex more favorable. (C) Model summarizing structural transition of RNA 4WJ tested here based on previous analysis [80]. The model shows that RNA 4WJs transitions between five conformations - four "closed" and one "open.". The observed small complex likely corresponds to binding to open conformation while the large complex to closed ones. (D) HMGB1a binds well to RNA 4WJ, however, instead of forming single large complex like MAEL HMG-box, it binds progressively, forming multiple intermediate complexes.

**Figure S5. Selection and structure determination of long RNA.**

(A) Multiple sequence alignment of regions identified through analysis of MAEL RIP-Seq datasets. Sequences correspond to L1_Md_F2 retrotransposon, are located on different chromosomes, and are highly conserved. Colored by percent identity (JalView). (B) Traces for individual regions after manual refinement of the alignment showing genomic locations and corresponding read coverage in each examined dataset. (C) Single, thermodynamically stable secondary structures generated in Mfold provided with covariance information for each identified genomic region. Shown are dG values of each structure as calculated by Mfold.

## Tables

**Table S1: Cloning oligoes**

**Table S2: DNA substrate sequences**

**Table S3: RNA substrate sequences**

**Table S4: PCR and IVT transcription oligoes**

**Table S5: Long RNA sequences**

**Table S6: Mfold constrains**

